# Ribosome remodeling under glucotoxic conditions impairs β-cell function

**DOI:** 10.64898/2026.11.11.765979

**Authors:** Abigael Cheruiyot Harvey, Jose Santos Navarro Arriola, Olasubomi Akintola, Ly Nguyen, Jennifer Hollister-Lock, Kanako Iwasaki, Susan Bonner-Weir, Cristina Aguayo-Mazzucato, Jean E. Schaffer

## Abstract

Diabetes develops when pancreatic β-cells fail to secrete sufficient insulin to meet metabolic demand. Chronic hyperglycemia accelerates β-cell failure and dysregulates mRNA translation before the onset of endoplasmic reticulum stress, but the underlying mechanisms remain unknown. Here, quantitative mass spectrometry of translating ribosomes isolated from β-cells under control and glucotoxic conditions revealed that sustained high glucose remodels ribosome composition by selectively decreasing Ribosomal Protein Lateral Stalk Subunit P1 (RPLP1) on actively translating ribosomes. Knockdown of RPLP1 in MIN6 cells and in primary islets reduced translation of insulin and impaired glucose-stimulated insulin secretion. Depletion of RPLP1 *in vivo* led to glucose intolerance. Together, our findings indicate that glucotoxicity remodels β-cell ribosomes, leading to selective mRNA translation impairment and decreased insulin secretion. This previously unrecognized mechanism linking metabolic overload to β-cell dysfunction suggests that preserving RPLP1 association with ribosomes could represent a therapeutic strategy for maintaining β-cell function in diabetes.

**IN BRIEF:** Hyperglycemia dysregulates pancreatic β-cell mRNA-specific translation and impairs glucose-stimulated insulin secretion. Here, Cheruiyot Harvey et al. reveal that sustained high glucose alters the protein stoichiometry of β-cell ribosomes, leading to impaired insulin translation and β-cell function.

**HIGHLIGHTS:**

- Sustained high glucose causes RPLP1 dissociation from translating ribosomes of β-cells.
- Sustained high glucose decreases phosphorylation of RPLP1, which is critical for polysome association.
- RPLP1-deficient β-cells have impaired insulin mRNA translation, GSIS, and glucose homeostasis.

## INTRODUCTION

Pancreatic β-cells within islets of Langerhans regulate systemic glucose homeostasis through glucose-stimulated insulin secretion (GSIS). Insufficient insulin secretion to meet metabolic demand is the defining feature of all forms of diabetes and results from loss or dysfunction of β-cells, impaired insulin action, or a combination of these processes.^1^ The resulting hyperglycemia plays a central role in initiation and propagation of cellular abnormalities underlying diabetic complications.^2,3^ Moreover, the observation that tight glycemic control improves β-cell function in both type 1 and type 2 diabetes indicates that hyperglycemia contributes to dysfunction and/or demise of the β-cells themselves.^4,5^ The metabolic overload of chronic hyperglycemia initiates mitochondrial dysfunction, oxidative stress, and endoplasmic reticulum (ER) stress that lead to loss of β-cell identity, dedifferentiation, and senescence.^2,6–9^ In experimental models, restoration of mitochondrial function and strategies that limit ER and oxidative stress have shown promise for preserving β-cell function.^10–12^ However, in humans with type 2 diabetes, approaches that reduce metabolic overload have the greatest impact on β-cell function.^13–15^ These limitations underscore the need to identify new targetable mechanisms that link metabolic overload to progressive β-cell failure.

The enormous secretory capacity of β-cells enables these cells to synthesize large quantities of insulin through a process that is rapidly and tightly regulated to respond to changing glucose levels. The insulin mRNA is highly abundant, comprising 30% of all β-cell mRNAs, and has a relatively long half-life,^16^ features that make this mRNA well-suited to translational regulation in response to the changing metabolic environment. Indeed, an acute, physiological rise in glucose stimulates not only insulin secretion, but also translation of the mRNAs for preproinsulin and proteins involved in glucose metabolism, prohormone processing, and insulin granule formation and release.^17^ In pathophysiological circumstances, prolonged exposure of β-cells to high glucose causes glucotoxicity, characterized initially by elevated basal insulin secretion and impaired GSIS and eventually leading to ER stress and depletion of insulin stores.^18,19^ Our recent work showed that sustained high glucose reduces translation of mRNAs for insulin and other proteins essential for uptake of glucose, its metabolism to produce ATP, and the formation and secretion of insulin granules.^20^ This translational dysregulation occurs prior to the onset of ER stress and without suppressing global protein synthesis, indicating that it is an early and selective defect in translation.

The classical model for protein synthesis posits that the ribosome performs core decoding and peptide synthesis reactions in a constitutive fashion, with output regulated by features of mRNAs, their binding proteins, and trans-acting factors.^21,22^ However, accumulating evidence challenges this view by showing eukaryotic ribosomes that differ in core ribosomal protein stoichiometry or rRNA modifications confer translational regulation of specific mRNAs. In embryonic stem cells and developing mouse embryos, ribosomes deficient in RPL10A, RPS25, or RPL38 preferentially translate distinct subsets of transcripts.^23,24^ In muscle and germ cells, ribosome protein paralogs that produce different translational outputs contribute to specialized cellular phenotypes.^25,26^ Heterogeneous ribosomes may regulate translation by impacting initiation, elongation, translation fidelity, termination efficiency, or co-translational folding.^26–30^

The contribution of ribosome heterogeneity to the specialization of pancreatic β-cell insulin secretion or its regulation is unknown. To address this knowledge gap, we performed quantitative mass spectrometry of translating ribosomes isolated from MIN6 β-cells cultured under control or glucotoxic conditions. We show that sustained high glucose causes dissociation of Ribosomal Protein Lateral Stalk Subunit P1 (RPLP1) from translating ribosomes in β-cells. Furthermore, we show that depletion of RPLP1 from β-cell ribosomes decreases translation of insulin, impairs GSIS, and compromises glucose tolerance *in vivo*. These findings provide evidence for dynamic ribosome remodeling in response to nutrient stress and identify changes in ribosome composition as a previously unrecognized mechanism for selective translational regulation in pancreatic β-cells.

## RESULTS

### Sustained high glucose decreases polysome-associated RPLP1 in MIN6 cells and islets

Sustained high glucose selectively decreases translation of mRNAs for critical proteins in GSIS, prior to induction of ER-stress or impairment of global translation.^20^ To understand mechanisms underlying this translational dysregulation, we examined the composition of actively translating ribosomes by quantitative proteomic analysis of polysomes isolated from MIN6 cells grown in media containing 25 mM (high) vs. 5.5 mM (low) glucose for 24 h (Figure 1A). Sustained high glucose decreased RPLP1 on polysomes by 50%, whereas abundance of other core ribosomal proteins was not significantly changed (Figure 1B, Supplemental Table 1). Immunoblots from independently generated MIN6 samples revealed no apparent change in total cellular abundance (input) of RPLP1, RPLP0 and RPL18A following 24 h incubation in high vs. low glucose media (Figure 1C). In the polysome fraction, sustained high glucose specifically decreased RPLP1 on the polysomes relative to other large subunit proteins. Endogenous RPLP1 migrated as a single band at the expected molecular weight (12 kDa) in whole cell lysates but consistently appeared as a doublet in precipitated protein from sucrose density gradient isolated polysomes (asterisk), potentially due to differences in post-translational modifications. These data indicate that stoichiometry of translating ribosomes was impacted by glucotoxic conditions.

**Figure. 1.**
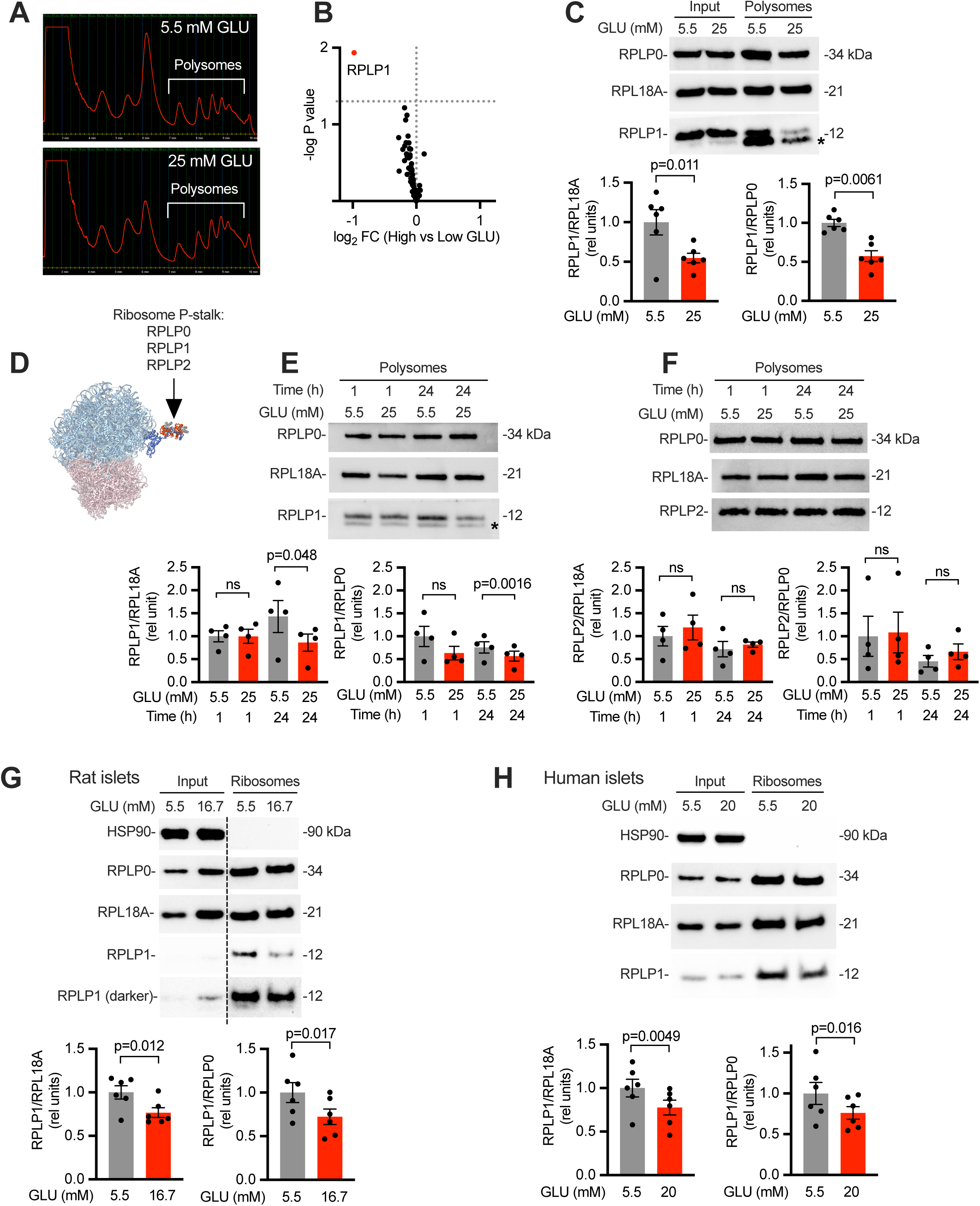
Sustained high glucose decreases RPLP1 association with translating ribosomes. **(A and B)** Sucrose density gradient fractionation of MIN6 cells cultured in media containing 5.5 vs. 25 mM glucose (GLU) for 24 h (**A**). Core ribosomal proteins in translating ribosomes detected using TMT-labeled proteomics from n = 4 independent experiments (**B**). **(C)** Representative polysome immunoblots following treatment of MIN6 cells for 24 h with 5.5 vs. 25 mM GLU. Quantification of RPLP1 in translating ribosomes relative to RPL18A or RPLP0 from n = 6 independent experiments. Asterisk indicates doublet observed for RPLP1 on immunoblots of precipitated, sucrose density gradient isolated polysomes. **(D)** Schematic of murine 80S ribosome cryo-EM structure (PDB: 9KDT) with the murine P-stalk complex (RPLP0, RPLP1, and RPLP2) modeled using AlphaFold 3. Only regions of high confidence prediction are shown for RPLP0 (amino acids 1-269), RPLP1 (amino acids 1-67), and RPLP2 (amino acids 1-64), because highly disordered carboxyl-terminal residues of these proteins are poorly resolved. **(E-F)** Representative polysome immunoblots following treatment of MIN6 cells for 1 h vs. 24 h with 5.5 vs. 25 mM GLU. Quantification of RPLP1 (**E**) and RPLP2 (**F**) relative to RPL18A or RPLP0 from n = 4 independent experiments. **(G)** Representative polysome immunoblots from rat islets treated with 5.5 vs. 16.7 mM GLU for 4 days. Quantification of RPLP1 relative to RPL18A or RPLP0 in translating ribosomes from n = 6 independent isolations. **(H)** Representative immunoblots from human cadaveric islets treated with 5.5 vs. 20 mM GLU for 48 h. Quantification of RPLP1 relative to RPL18A or RPLP0 in translating ribosomes from n = 6 donors. Means ± standard error (SE). P-values by two-tailed paired t-tests. ns, not significant

RPLP1 is a constituent of the flexible ribosome P-stalk, part of the GTPase-associated center on the large subunit that functions in the docking and activation of translational GTPases and activation of the stress kinase GCN2.^31–33^ The stalk is a pentameric structure consisting of one copy of RPLP0 and 2 copies each of RPLP1 and RPLP2 (Figure 1D).^34^ Decrease in polysome-associated RPLP1 was observed following high glucose treatment for 24 h, which impairs β-cell function,^20^ but not with acute glucose stimulation for 1 h, which increases GSIS and translation of mRNAs required for insulin production and secretion (Figure 1E).^17^ Notably, RPLP2 abundance on translating polysomes was unchanged by sustained high glucose exposure (Figure 1F). Thus, glucotoxic conditions specifically decreased RPLP1 association with translating ribosomes in MIN6 cells.

MIN6 cells are a useful insulinoma-derived proliferating β-cell model that supports GSIS and provides sufficient material to perform high-throughput discovery studies, such as quantitative proteomic analysis of polysome fractions.^35^ To determine whether glucotoxicity also decreases RPLP1 on translating ribosomes in primary islets, we treated rat and human islets with sustained high glucose conditions previously shown to impair GSIS and alter mRNA-specific translation without inducing ER stress or impairing global protein synthesis rates.^20^ Consistent with observations in MIN6 cells, translating ribosome-associated RPLP1 was reduced in rat islets exposed to 16.7 mM glucose vs. 5.5 mM glucose for four days (Figure 1G). Similarly in human islets, we observed decreased RPLP1 association with translating ribosomes following treatment for two days with 20 mM glucose vs. 5.5 mM glucose (Figure 1H). These findings establish that reduced polysome-associated RPLP1 is a conserved response to sustained high glucose in MIN6 cells and primary islets.

### Decrease in polysome RPLP1 reflects dissociation of RPLP1 from ribosomes

RPLP1 and RPLP2 are unusual, acidic ribosomal proteins that also have stable free cytoplasmic pools and exchange between ribosome-bound and soluble states.^36–38^ Given that total lysate RPLP1 was unchanged during glucotoxic stress, we hypothesized that the decrease in polysome RPLP1 reflected movement from ribosome-bound to soluble pools. However, using commercially available rabbit polyclonal antibody to RPLP1, we were unable to detect soluble RPLP1. To enhance sensitivity of detection, we used lentiviral transduction to express RPLP1 with an amino-terminal FLAG tag (Figure 2A). The 18 kDa FLAG-tagged protein was expressed at low levels compared to the 12 kDa endogenous RPLP1 doublet (Figure 2B, triangle denotes FLAG-tagged RPLP1). Nonetheless, the tagged protein exhibited similar depletion from polysomes following sustained high glucose treatment in MIN6 (Figures 2C and 2D). Using a monoclonal α-FLAG antibody to immunoprecipitate the epitope-tagged protein, we observed that the decrease in polysome-associated FLAG-RPLP1 was accompanied by a 4-fold increase in the small soluble pool of FLAG-RPLP1 (Figure 2E).

**Figure 2.**
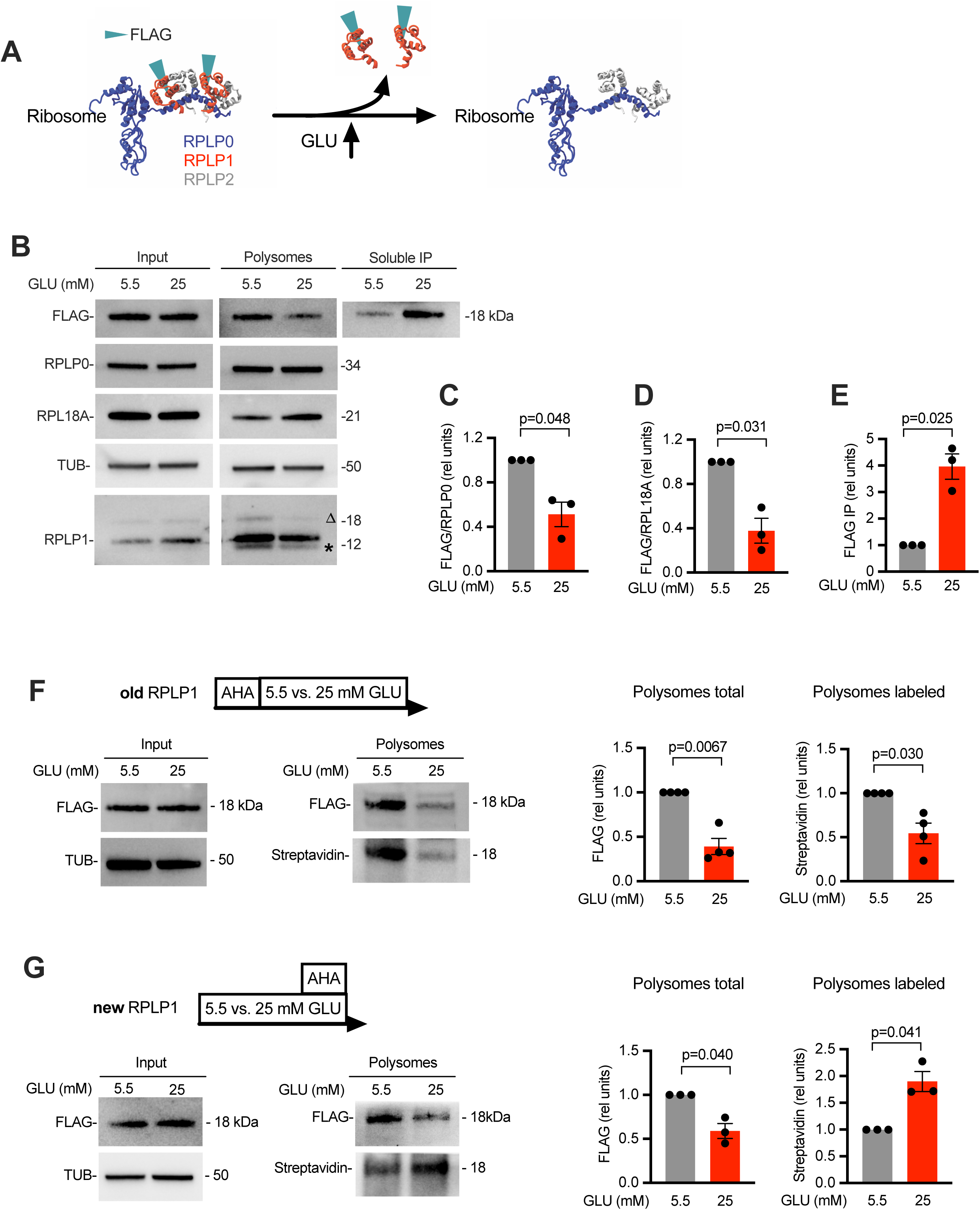
Decrease in polysome RPLP1 reflects dissociation of protein from existing ribosomes. **(A)** Schematic of FLAG-tagged murine RPLP1 (FLAG-RPLP1) in complex with RPLP0 and RPLP2, as predicted by AlphaFold 3. Regions of high confidence prediction are shown for RPLP0 (amino acids 1-269), RPLP1 (amino acids 1-67), and RPLP2 (amino acids 1-64). **(B-E)** Input, polysomes, and soluble fractions isolated from MIN6 cells expressing FLAG-RPLP1 treated with 5.5 vs. 25 mM GLU for 24 h. Representative immunoblots of input and polysomes using α-FLAG, α-RPLP0, α-RPL18A, α-tubulin (TUB) or polyclonal α-RPLP1 antibodies for detection and representative immunoblot of FLAG-RPLP1 immunoprecipitated from soluble fraction with monoclonal α-FLAG (**B**). Asterisk on α-RPLP1 blot indicates doublet from endogenous protein; triangle indicates FLAG-RPLP1. Quantification of polysome FLAG-RPLP1 relative to RPLP0 (**C**) and RPL18A (**D**), and quantification of FLAG-RPLP1 immunoprecipitated from soluble fraction (**E**). **(F-G)** Metabolic labeling of pre-existing (old, **F**) and newly synthesized (new, **G**) proteins with azidohomoalanine (AHA) with low and high glucose treatment as indicated in diagrams. Representative immunoblots of input and α-FLAG-immunoprecipitated polysomes with detection of total FLAG-RPLP1 by α-FLAG and metabolically labeled FLAG-RPLP1 by streptavidin-HRP. Quantification of polysome-associated total FLAG-RPLP1 (FLAG immunoblot) and polysome-associated metabolically-labeled RPLP1 (streptavidin blot) relative to input FLAG-RPLP1. Means of 3 to 4 independent experiments ± SE. P-values by one-sample t-test and Wilcoxon test.

Two potential mechanisms could explain co-incident decrease in polysome-associated protein and increase in soluble protein without change in total protein. Glucotoxicity might induce dissociation of RPLP1 from pre-existing, mature ribosomes. Alternatively, newly synthesized RPLP1 may fail to associate with newly synthesized ribosomes. To distinguish between these possibilities, we performed metabolic labeling studies, in which cells were incubated with the clickable methionine analogue azidohomoalanine (AHA); polysomes were isolated; biotin was added by click addition; FLAG-tagged RPLP1 was immunoprecipitated; and recovered protein was analyzed by FLAG immunoblotting for total RPLP1 and streptavidin blotting for newly synthesized RPLP1. First, we labeled preexisting ribosomes using an 8 h incubation with AHA, followed by removal of AHA from the media during treatment for 24 h with high vs. low glucose media. Sustained high glucose decreased pre-labeled (previously synthesized) FLAG-RPLP1 on the polysomes to a similar extent as the decrease in immunodetectable total FLAG-RPLP1 (Figure 2F). Second, we labeled proteins synthesized during the final 8 h of high vs. low glucose media treatment. Although sustained high glucose decreased immunodetectable FLAG-RPLP1 on polysomes, AHA-labeled (newly synthesized) RPLP1 significantly increased on polysomes (Figure 2G). With both labeling approaches, AHA-labeled RPLP1 was not detected in the soluble fraction, likely due to inefficiencies inherent in metabolic labeling. Together, our data support a model in which glucotoxic conditions cause dissociation of RPLP1 from the ribosome.

### Phosphorylation on carboxyl-terminal serines mediates glucotoxic effects on RPLP1 localization

In *Saccharomyces cerevisiae*, phosphorylation of the acidic exchangeable ribosomal proteins promotes affinity for the ribosome, and the cytoplasmic pools of these proteins are unphosphorylated.^39,40^ Furthermore, phosphoproteomic analysis in MIN6 cells detected phosphorylation of RPLP1 on serine residues 101 and 104.^41^ To determine whether exogenously expressed FLAG-RPLP1 is phosphorylated in MIN6 cells, we analyzed immunoprecipitates from cells transduced with FLAG-RPLP1 by immunoblotting and staining with Pro-Q Diamond for phosphoproteins. The robust signal by Pro-Q Diamond staining at the expected molecular weight of FLAG-RPLP1 was abrogated by pre-treatment of the lysate with alkaline phosphatase (Figure 3A). These data indicate that epitope-tagged RPLP1 is phosphorylated in MIN6 cells.

**Figure 3.**
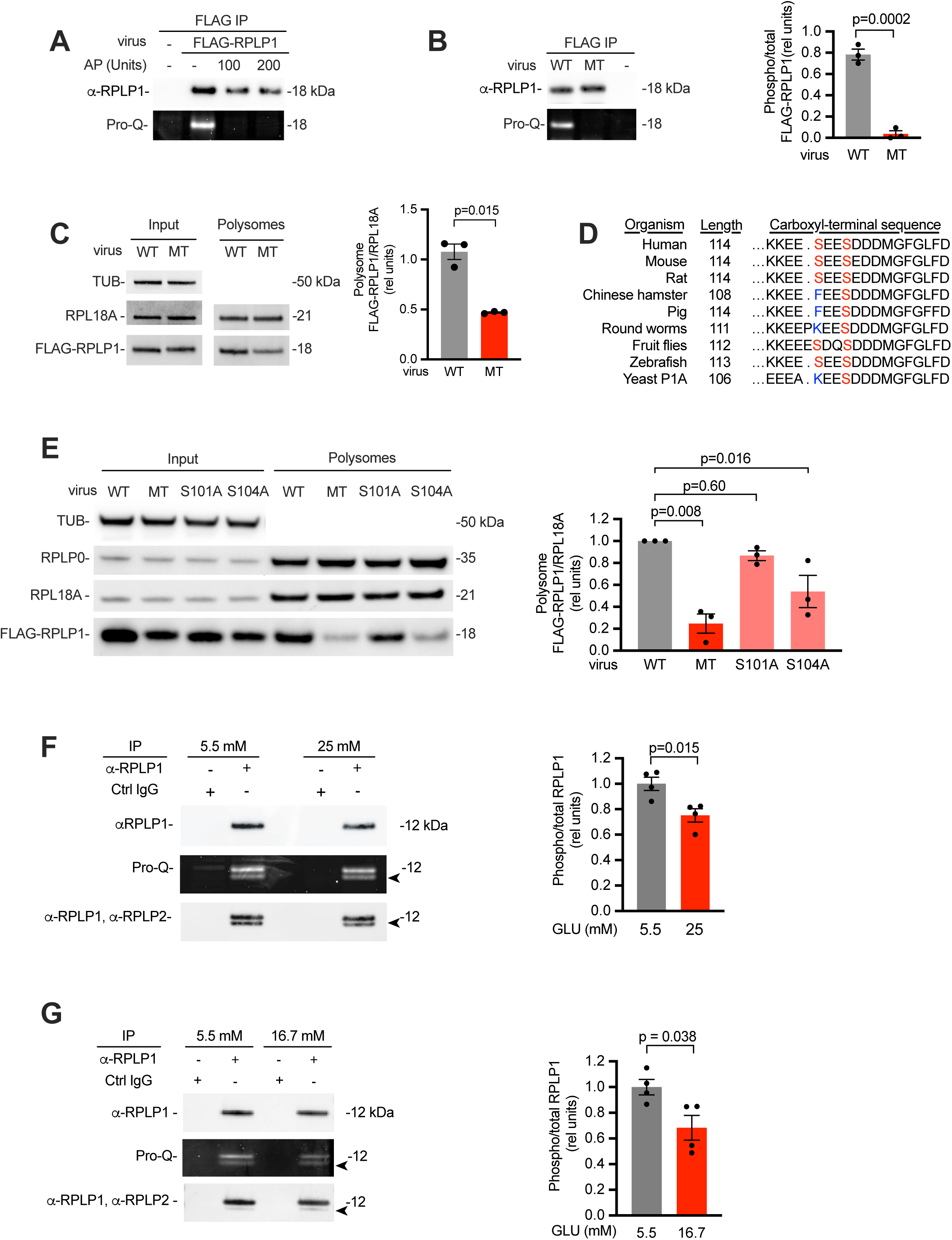
RPLP1 phosphorylation is modulated by sustained high glucose. **(A)** Immunoprecipitated FLAG-RPLP1 from MIN6 cells analyzed by SDS-PAGE. Representative immunoblot and Pro-Q Diamond detection without and with alkaline phosphatase (AP) treatment. **(B)** Wild type (WT) and S101A/S104A mutant (MT) FLAG-RPLP1 immunoprecipitated from MIN6 cells and analyzed by SDS-PAGE. Representative immunoblot and Pro-Q Diamond detection. Quantification of phosphorylated FLAG-RPLP1 relative to immunoprecipitated FLAG-RPLP1. **(C)** Polysome analysis of MIN6 cells expressing WT or MT FLAG-RPLP1. Representative immunoblots. Quantification of polysome FLAG-RPLP1 (WT or MT) relative to RPL18A and normalized to input FLAG-RPLP1 (WT or MT) relative to tubulin (TUB). **(D)** Evolutionary conservation across species of RPLP1 carboxyl-terminal residues. Amino acid residues corresponding to serine 101 and 104 of the mouse sequence in red or blue. (E) Representative immunoblot of input and polysomes from cells expressing WT, MT, S101A FLAG-RPLP1 or S104A FLAG-RPLP1. Quantification of polysome FLAG-RPLP1 (WT or mutant) relative to RPL18A and normalized to input FLAG-RPLP1 (WT or MT) relative to tubulin (TUB). (**F-G**) Representative RPLP1 immunoblot and Pro-Q Diamond detection of immuno-precipitated endogenous RPLP1 from MIN6 cells treated with 5.5 vs 25 mM GLU for 24 h (**F**) or from rat islets treated with 5.5 vs. 16.7 mM GLU for 4 days (**G**). RPLP1 immunoblots were re-stained with α-RPLP2, demonstrating the lower band on Pro-Q Diamond staining is co-immunoprecipitating RPLP2 (arrowheads). Quantification of phospho-RPLP1 (upper band only) relative to immunodetected RPLP1. Means ± SE for 3 to 4 independent experiments. P-values by unpaired two-tailed t-tests (**B, C, F, G**) or one-way ANOVA (**E**).

To evaluate the sites of RPLP1 phosphorylation, we generated a construct in which serines 101 and 104 of the endogenous RPLP1 sequence were substituted with alanine to prevent phosphorylation. Although wild type FLAG-RPLP1 (WT) and FLAG-RPLP1-S101A/S104A (MT) were expressed at comparable levels, mutation of residues 101 and 104 led to complete loss of phosphorylation signal compared to WT (Figure 3B). We then asked how the lack of phosphorylation of the MT impacted polysome association. Immunoblots of sucrose density gradient fractions from cells expressing either WT or MT demonstrated comparable expression of FLAG-tagged proteins in inputs (Figure 3C). However, mutation of serine 101 and serine 104 to alanine significantly reduced polysome association compared to wild type RPLP1. Thus, serines 101 and 104 collectively account for the detectable phosphorylation signal of FLAG-RPLP1 and their substitution to alanine reduces polysome association.

RPLP1 serine residues at positions 101 and 104 lie within the evolutionarily conserved carboxyl terminus of the protein. While some substitutions at serine 101 occur in RPLP1 orthologs across species, serine 104 is invariant, raising the possibility that it could play a dominant role in RPLP1 polysome association (Figure 3D). To assess the individual contributions of these residues, we compared localization of FLAG-RPLP1 single mutants S101A or S104A to WT and MT. Mutation of serine 104 to alanine significantly decreased polysome association, whereas mutation of serine 101 alone only trended to decrease association (Figure 3E). MT exhibited the largest reduction in polysome association. These findings indicate that S104 is an important determinant of polysome association and suggest that S101 contributes to a lesser extent.

Previous analyses of the MIN6 phosphoproteome demonstrated no significant change in phosphorylation of RPLP1 following acute treatment with high glucose,^41^ conditions under which we do not observe dissociation of RPLP1 from the polysomes (Figure 1E). To determine whether glucotoxic conditions that prompt dissociation of endogenous RPLP1 from polysomes are associated with dephosphorylation of the protein, we analyzed endogenous RPLP1. Following sustained high glucose, immunoprecipitated RPLP1 was analyzed by immunoblotting and Pro-Q Diamond staining. Sustained high glucose decreased RPLP1 phosphorylation in MIN6 cells by 25% (Figure 3F) and in rat pancreatic islets phosphorylation was decreased by 32% (Figure 3G). Collectively, our data are consistent with a model in which sustained high glucose decreases association of RPLP1 with translating ribosomes by decreasing phosphorylation of RPLP1.

### RPLP1 knockdown dysregulates mRNA-specific translation and GSIS

Although RPLP1 is not essential for protein synthesis,^42^ presence or absence of RPLP1 on ribosomes has been shown to regulate mRNA-specific translation in mouse embryonic stem cells and human melanoma cells.^43,44^ To determine the functional consequences of decreased RPLP1 on translating ribosomes in β-cells, we knocked down RPLP1 in MIN6 cells using lentiviral short hairpin RNAs (shRNAs) targeting RPLP1. We achieved 40% knockdown with two independent shRNAs (Figure 4A). Knockdown had no significant impact on global protein synthesis, as measured by incorporation of the puromycin analog, O-propargyl-puromycin (OPP) (Figure 4B). However, knockdown significantly decreased translation efficiency of mRNAs known to be critical for GSIS, including *Ins1*, *Pfkfb3*, *Slc2a2*, *Slc30a8*, and *Igf2* (Figure 4C). There was a trend for decrease in *Ins*2 translation efficiency that did not reach significance. Furthermore, RPLP1 knockdown decreased GSIS in MIN6, with an average 67% decrease in stimulation index (Figures 4D and 4E). Together, these results indicate that RPLP1 plays an important role in mRNA-specific translation and GSIS in a model of β-cells.

**Figure 4.**
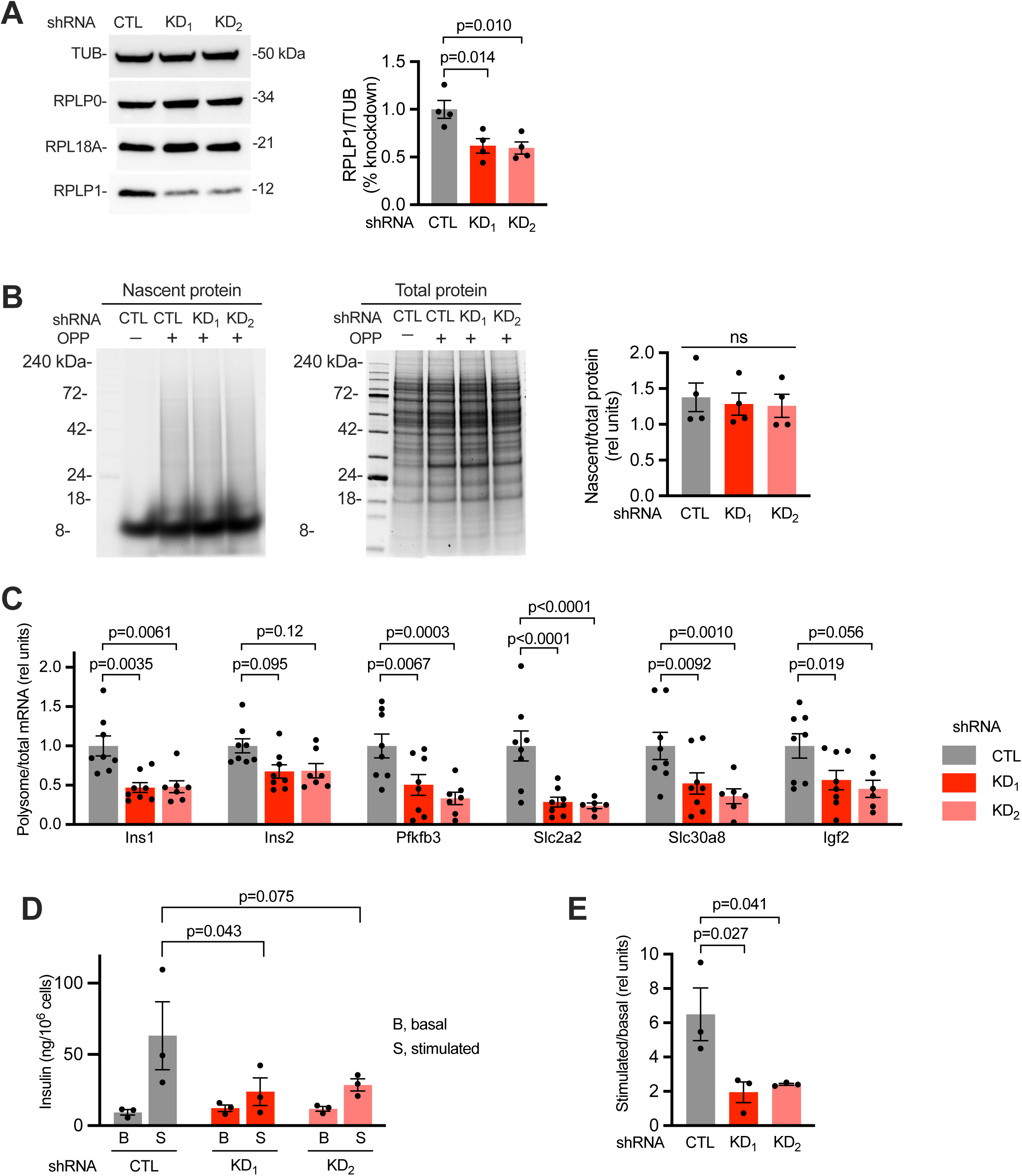
RPLP1 knockdown dysregulates mRNA-specific translation and GSIS in MIN6 cells. (A) Representative immunoblots of MIN6 cells transduced with control (CTL) or RPLP1-targeting (KD_1_, KD_2_) shRNA lentivirus. Quantification of RPLP1 relative to tubulin. RPLP0 and RPL18A as controls. **(B)** CTL and KD cells incubated with O-propargyl-puromycin (OPP) for 1 h. SDS-PAGE of nascent (CLICK-Alexafluor 647) and total (Coomassie stain) proteins with quantification of nascent/total protein. **(C)** Translation efficiency of *Ins1*, *Ins2, Pfkfb3*, *Slc2a2*, *Slc30a8*, and *Igf2* by qPCR quantification of polysome-associated/total RNA. **(D and E)** Insulin secretion in GSIS analysis (**D**) with quantification of stimulation index (**E**). Means of 3 to 8 independent experiments ± SE. P-values by one-way (**A, B, E**) or two-way (**C, D**) ANOVA.

We also used lentiviral shRNAs to examine the impact of RPLP1 knockdown in primary rat islets. By transducing freshly isolated islets in which vascular channels are most likely to remain open,^45^ we achieved an average of 54% decrease in islet RPLP1 (Figure 5A). Knockdown decreased translation efficiency for *Ins1* and *Ins2* mRNAs, although it did not affect translation efficiency of *Pfkfb3, Slc2a2, Slc30a8, or Igf2* (Figure 5B). The more limited effect on translation in islets compared to MIN6 cells may reflect differences between a clonal β-cell-derived line and primary islets which include α-, δ-, PP, and ε-cells in addition to β-cells. Nonetheless, RPLP1 knockdown tended to increase basal secretion of insulin, and it decreased stimulation index (Figure 5C and 5D), mirroring the impact of sustained high glucose observed previously in islets and consistent with findings in senescent and failing β-cells.^20,46^ Thus, presence of RPLP1 in pancreatic islets regulates mRNA-specific translation and is critical for β-cell function.

**Figure 5.**
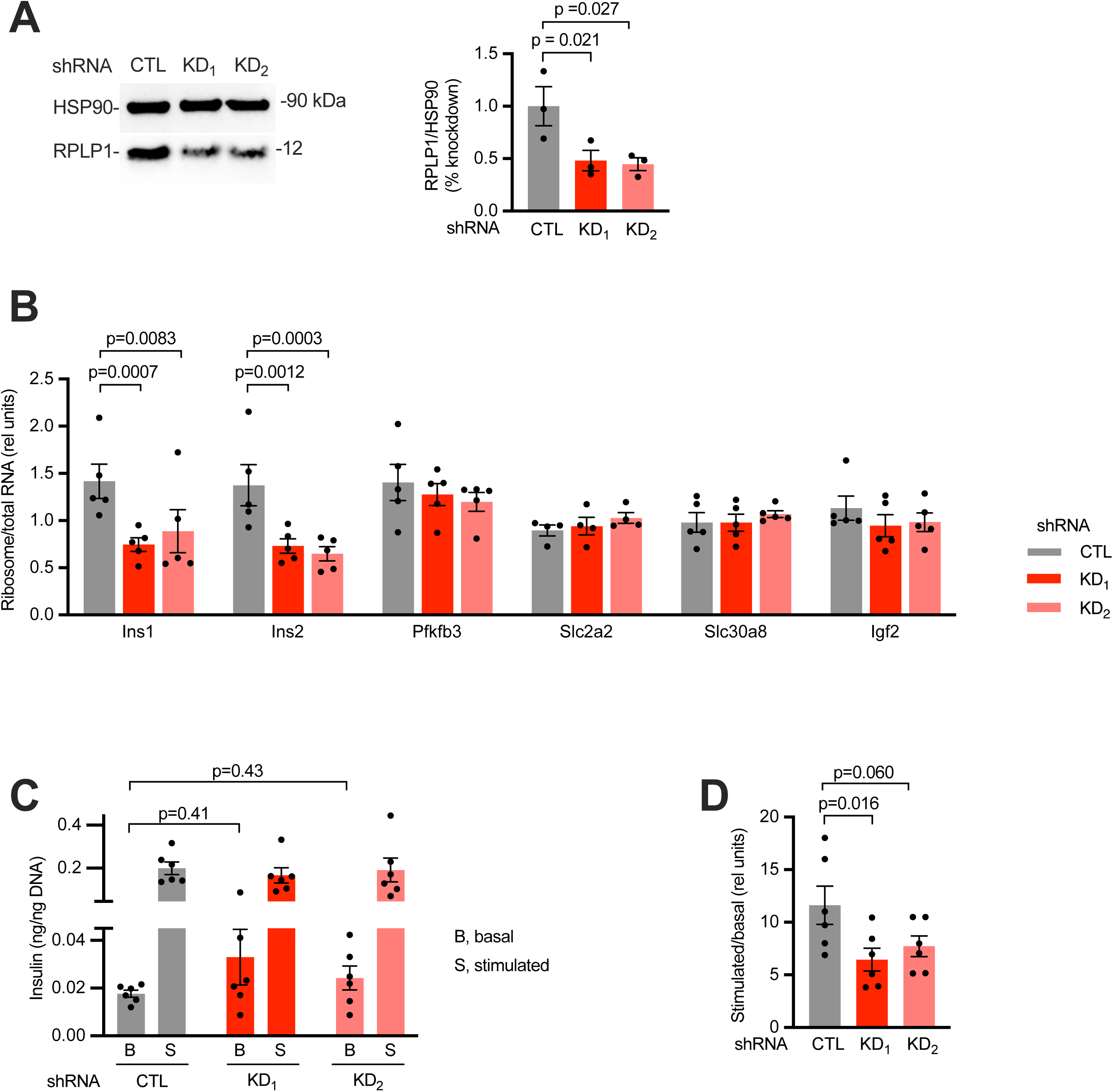
RPLP1 knockdown dysregulates rat islet insulin mRNA translation and GSIS. **(A)** Representative immunoblots of rat islets transduced with control (CTL) or RPLP1-targeting (KD_1_, KD_2_) shRNA lentivirus. Quantification of RPLP1 normalized to HSP90. **(B)** Translation efficiency of *Ins1*, *Ins2*, *Pfkfb3*, *Slc2a2*, *Slc30a8*, and *Igf2* determined as ribosome-bound mRNA relative to total mRNA in CTL, KD_1_ and KD_2_ islets. **(C and D)** GSIS analysis (**C**) and quantification of stimulation index **(D)**. Means of 3 to 6 independent experiments ± SE. P-values by one-way (**A, C** (basal glucose only), **D**) or two-way (**B**) ANOVA.

### β-cell RPLP1 is critical for glucose homeostasis *in vivo*

To extend these findings to physiological models, we first tested whether knockdown of RPLP1 impacts the ability of transplanted islets to support glucose homeostasis (Figure 6A). Immunocompromised NCG mice were treated with streptozotocin (STZ) to target β-cells and induce a diabetic-like state. Fed blood glucose in NCG recipient animals averaged 450 mg/dl prior to islet transplantation (Figure 6B). Isolated rat islets were transduced with RPLP1-targeting shRNA or control scrambled shRNA, achieving 53% stable knockdown of *Rplp1* mRNA without affecting insulin content (Figures 6C and 6D). RPLP1 knockdown and control transduced islets (400 islet equivalents [IEQ]) were transplanted into diabetic recipients, which were followed for two weeks prior to analysis. There were no differences in fed morning glucose (not shown) or fasting glucose between animals that received islets with RPLP1 knockdown vs. those that received control scrambled shRNA islets (Figure 6E). However, animals transplanted with islets with RPLP1 knockdown had increased glucose area under the curve during intraperitoneal glucose tolerance testing (Figures 6F and 6G). These findings indicate that decreased RPLP1 expression in islets diminished their ability to support glucose homeostasis.

**Figure 6.**
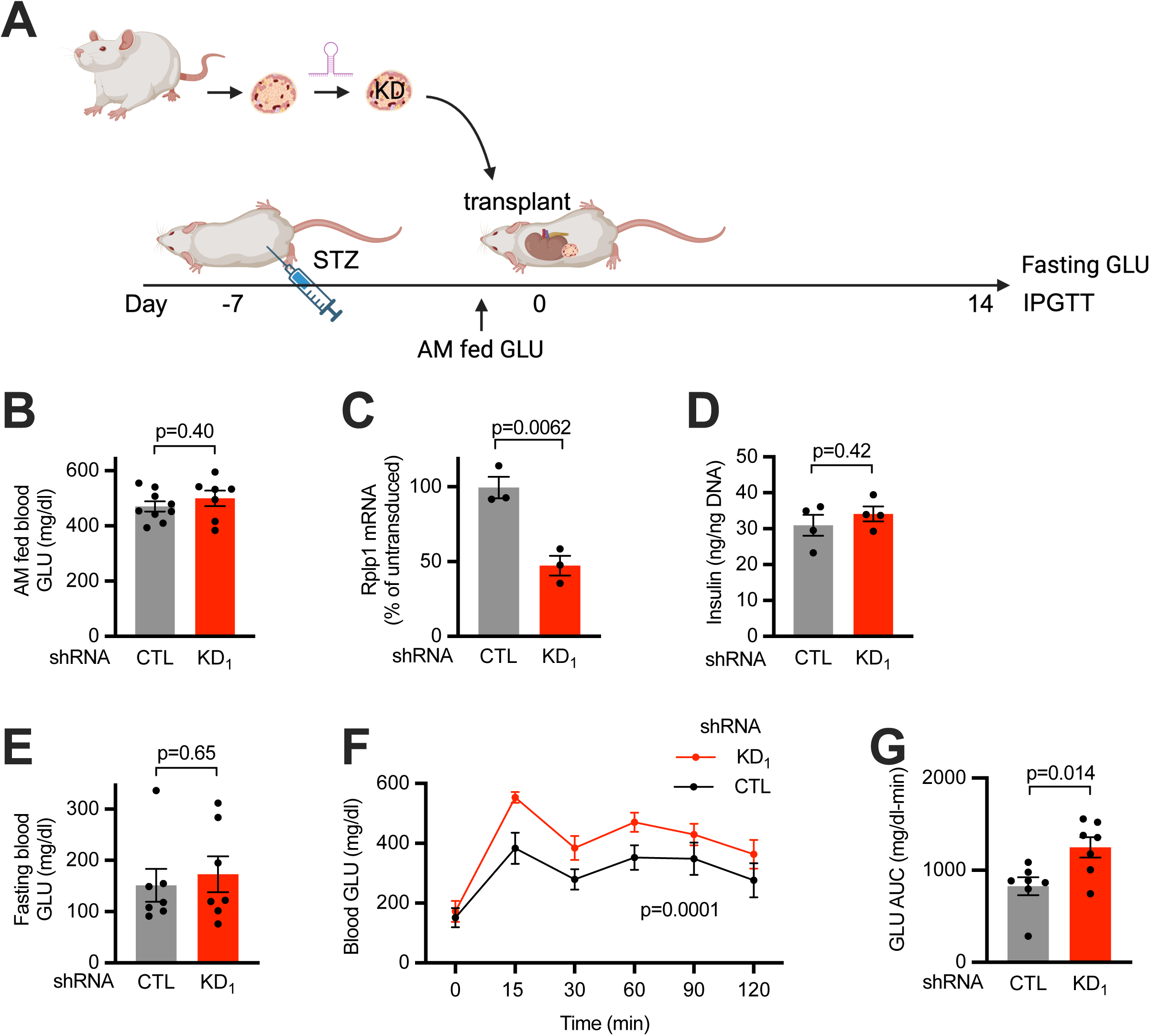
RPLP1 knockdown decreases function of transplanted islets *in vivo*. **(A)** Schematic showing rat islet transplantation experimental design. **(B-D)** Analyses prior to transplant. Morning fed blood glucose of NCG recipient mice 7 days after injection with streptozotocin (STZ) and prior to transplant (**B**). n = 7 to 9 mice per group. Quantification of *Rplp1* mRNA (**C**) and insulin content (**D**) from rat islets transduced with CTL or KD_1_ lentiviral shRNA. n = 3 to 4 independent isolations. (**E-G**) Analyses following transplant. Fasting blood glucose 14 days following islet transplantation (**E**). N = 7 per group. Intraperitoneal glucose tolerance testing (IPGTT, **F**) and calculated glucose area under the curve (AUC, **G**) for n = 7 mice per group. Means ± SE. P-values by two-tailed t-test (**B-E, G**) or two-way ANOVA (**F**).

In a second *in vivo* model, we used gene editing to test the contribution of RPLP1 expression specifically in pancreatic β-cells to systemic glucose homeostasis (Figure 7A). Mice expressing insulin-promoter driven Cas9 were injected with AAV8 for expression gRNAs targeting *Rplp1* or control scrambled gRNA. Morning fed glucose was higher in mice with β-cell *Rplp1* editing, but fasting blood glucose was not significantly different (Figures 7B and 7C), findings consistent with other mouse models of early β-cell dysfunction in which fed glucose reveals mild to moderate β-cell dysfunction, even when fasting glucose remains normal.^47^ Glucose tolerance testing demonstrated impaired glucose clearance in mice with *Rplp1* editing and increased glucose area under the curve compared to animals that received control gRNA (Figures 7D and 7E). Immunostained pancreas sections revealed decreased islet RPLP1 staining in animals that received *Rplp1*-targeting gRNAs (Figures 7F-7G). Decreased intensity of islet insulin staining in these animals suggested insulin content was also decreased (Figure 7H). Consistent with these findings, isolated islets from animals that received *Rplp1* gRNAs showed decreased RPLP1 by immunoblotting (Figure 7I) and decreased insulin content (Figure 7J). Together, these data indicate that β-cell expression of RPLP1 is critical for maintaining β-cell function and systemic glucose homeostasis.

**Figure 7.**
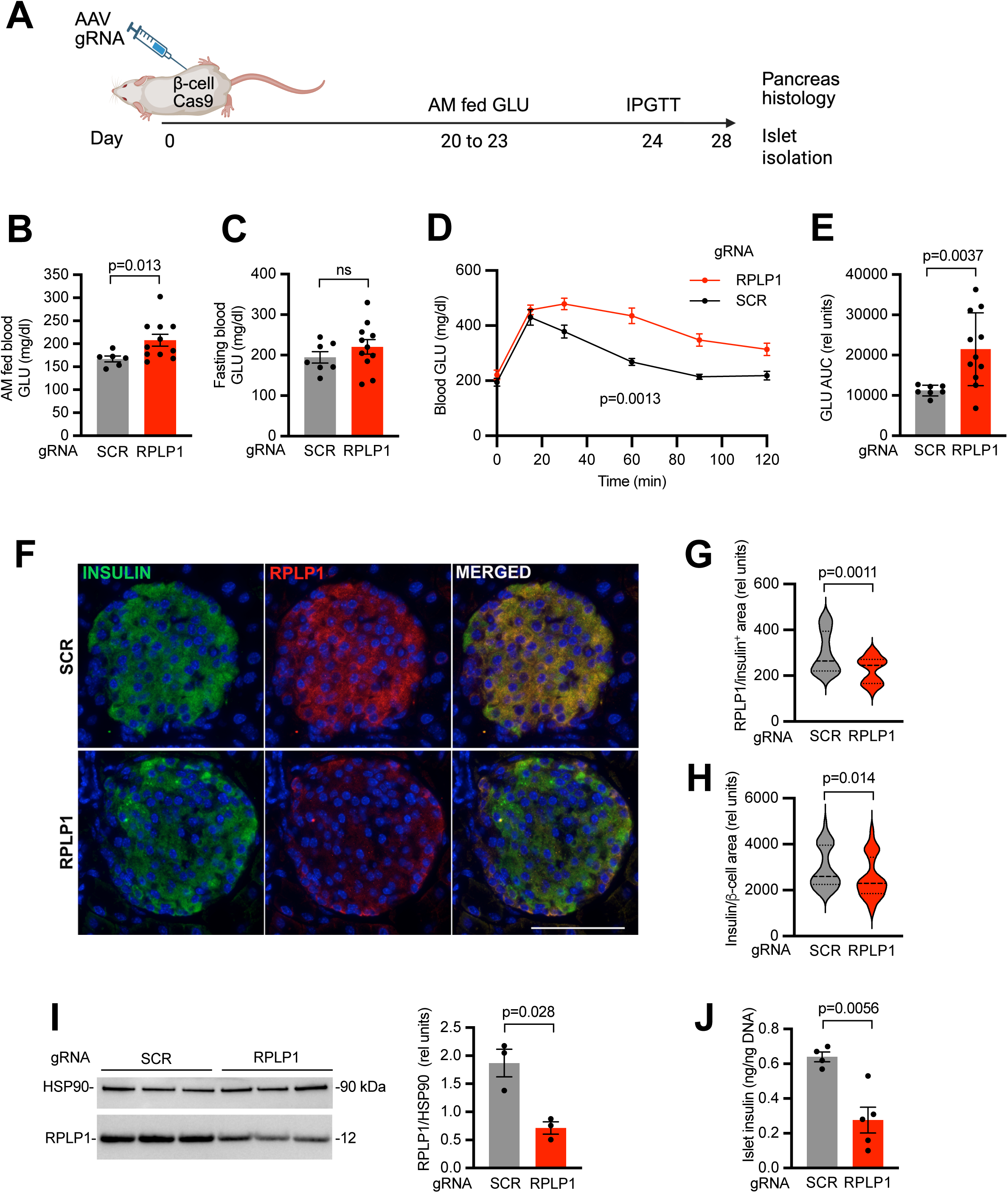
RPLP1 editing in β-cells impairs glucose homeostasis. **(A)** Schematic showing the *in vivo* editing experimental design. **(B-E)** Measures of glucose homeostasis for RPLP1 (n = 11) and SCR (n = 7) gRNA treated mice. Morning fed blood GLU quantified between days 20 and 23 (**B**). Fasting blood GLU on day 24 at the start of IPGTT (**C**). IPGTT (**D**) and calculated glucose area under the curve (**E**). **(F-H)** Representative immunofluorescent staining of pancreas sections for RPLP1 (red), insulin (green), and DAPI (blue) (**F**). Scale bar 70 µm. Quantification of RPLP1/insulin^+^ area (**G**) and intensity of insulin staining/β-cell area (**H**). **(I)** Representative immunoblot analysis of islets isolated at day 28 after AAV. Quantification of RPLP1 relative to HSP90. **(J)** Quantification of insulin content of islets isolated at day 28 after AAV. Means ± SE. P-values by two-tailed unpaired t-tests (**B, C, E, I, J**), Mann-Whitney two-tailed t-test (**G, H**), or by 2-way ANOVA (**D**).

## DISCUSSION

Here we show that the core protein stoichiometry of actively translating β-cells ribosomes is modified by glucotoxic conditions. Exposure to sustained high glucose specifically reduced RPLP1 association with translating ribosomes in MIN6 insulinoma cells and in rat and human islets. This ribosome remodeling reflected dissociation of RPLP1 from pre-existing ribosomes to a cytosolic pool and was associated with dephosphorylation. In MIN6 cells and islets, knockdown of RPLP1 impaired mRNA-specific translational regulation and decreased GSIS. Transplantation of islets with RPLP1 knockdown into diabetic mice and Cas9-editing of the endogenous *Rplp1* locus in β-cells led to impaired glucose tolerance. Together, our findings establish that β-cell ribosomes are dynamically remodeled in response to the nutrient environment in ways that dysregulate mRNA-specific translation and compromise β-cell function.

RPLP1 is one of three acidic ribosomal proteins that form a pentameric structure on the flexible P-stalk of the 60S ribosomal subunit. One molecule of RPLP0 and 2 molecules each of RPLP1 and RPLP2 are added to the periphery of the ribosome late during ribosome biogenesis, when the nascent structure reaches the cytoplasm.^48^ Unlike most ribosomal proteins that are rapidly degraded if not incorporated into the ribosome structure,^49^ RPLP1 and RPLP2 have been shown to exist in stable cytosolic pools that exchange with ribosome-bound forms of these proteins.^37^ Phosphorylation of the carboxyl-terminal sequences of RPLP1 and RPLP2 drives ribosome association of these P-stalk proteins.^50–53^ Our analysis of RPLP1 mutants established the contributions of serine residues 101 and 104 to both RPLP1 phosphorylation and polysome association in mammalian cells and identifies a new target of altered pancreatic β-cell signaling under metabolic stress conditions. RPLP1 has been shown to be phosphorylated by casein kinase 2 (CK2), a constitutively active protein kinase.^54^ Our finding that decreased ribosome-associated and concomitant increased soluble RPLP1 resulted from dissociation of the protein from the ribosome supports a model in which sustained high glucose induces expression and/or activity of a phosphatase that acts on RPLP1. Identification of the putative phosphatase and determination of whether it can be specifically targeted will be the subject of future studies.

RPLP1 and RPLP2 form heterodimers that bind to distinct sites on the carboxyl terminus of RPLP0.^55^ Although glucotoxicity caused RPLP1 to dissociate from translating ribosomes, we observed no change in polysomal content of RPLP2. This is consistent with a previously published study showing that RPLP2 associates with translating ribosomes in mammalian cells even in RPLP1 knockout cells.^56^ Despite their highly conserved carboxyl-terminal sequences,^57^ RPLP1 and RPLP2 share only 35% overall amino acid sequence identity in humans and mice. Divergent sequences of these proteins likely contribute to their distinct responses to glucotoxic stress.

As a component of the ribosome GTPase-activating center, the flexible ribosome P-stalk contributes to regulation of translation through recruitment and activation of translational GTPases and interactions with translation factors.^33^ During amino acid starvation, binding and activation of GCN2 that initiates the integrated stress response requires both RPLP1 and RPLP2.^31,32^ In vitro evidence also supports specific binding of both RPLP1 and RPLP2 to translation factor EEF2.^58^ Although knockdown or knockout of RPLP1 or RPLP2 has minimal impact on global translation, these proteins are required for translation of specific mRNAs in murine embryonic stem cells and immortalized human carcinoma cells.^56,59,60^ In a recent study, Ribo-Tweezer technology enabled rapid removal of RPLP1 from mature ribosomes of mouse embryonic stem cells, leading to translational upregulation and downregulation of specific mRNAs.^43^ In several independent RPLP1 loss-of-function studies, down-regulated mRNAs are enriched for proteins with membrane-spanning domains.^43,59^ Our knockdown studies demonstrate that reduced RPLP1 abundance in β-cells selectively affects the translation efficiency of mRNAs important for metabolism-coupled insulin secretion. This group included secreted, soluble, and membrane proteins. The basis for selection of specific mRNAs by ribosomes containing different compositions of stalk proteins is not well understood, in part because the P-stalk proteins are not well resolved in structural studies due to conformational flexibility of this region of the ribosome.^61,62^ Future studies will be required to define the features that render specific client mRNAs sensitive to the presence or absence of ribosome-associated RPLP1.

The specialized-ribosome hypothesis posits that ribosomes with distinct compositions can contribute to differences in translation specificity. Determining whether differences in protein composition, rRNA modifications or associated factors directly alter transcript selection, initiation, elongation, or termination has been challenging, because multiple simultaneous cellular mechanisms can regulate translation.^63,64^ Nonetheless, accumulating evidence supports the model that heterogeneous ribosomes co-exist within cells and contribute to different translational outputs, often in specialized cells or different developmental contexts.^23–25^ Ribosome protein composition and translational outputs are also acutely altered by osmotic stress and high pH in *S. cerevisiae* and by cytokine signaling in melanoma cells.^44,65^ Our finding that sustained high glucose leads to decreased presence of RPLP1 on translating ribosomes and dysregulates mRNA translation for critical GSIS proteins provides an example in which ribosome composition and function are dynamically regulated in a specialized cell type in response to the changing metabolic environment.

At the core of β-cells’ ability to produce sufficient insulin to maintain glucose homeostasis is their capacity to rapidly upregulate the translation of insulin and proteins required for coupling its secretion to metabolism.^17,66^ Selective β-cell translational regulation in response to acute increases in glucose has been linked to sequence and/or structural features of the mRNAs themselves and activity of translation initiation factors.^67–70^ Our *ex vivo* and *in vivo* studies suggest that RPLP1 is particularly important for maintaining translation efficiency of GSIS proteins. In both MIN6 cells and islets, RPLP1 knockdown diminished insulin stimulation index, by decreasing secretion in response to stimulatory glucose in MIN6 and by increasing basal secretion in islets. Knockdown or editing of *Rplp1* impaired glucose homeostasis *in vivo*. The finding that presence of RPLP1 on β-cell ribosomes is critical for maintenance of glucose homeostasis raises the possibility that interventions to prevent RPLP1 dissociation from ribosomes could preserve or restore β-cell function.

### Limitations of the study

The molecular events that determine RPLP1 dissociation from the ribosome remain incompletely understood. Although our Pro-Q Diamond staining studies indicate that sustained high glucose decreases phosphorylation of RPLP1, our efforts to obtain direct phosphoproteomic evidence of change in phosphorylation were limited by two factors. First, the carboxyl terminus of RPLP1 containing serine residues 101 and 104 is highly acidic. Peptides containing this region of the protein ionize inefficiently. Second, tandem lysine residues in the RPLP1 carboxyl terminus led to generation of multiple peptide species upon digestion, distributing the detectable signal across different peptides. To confirm our findings with Pro-Q Diamond staining, we tested several general anti-phosphoserine antibodies, but these failed to recognize phosphorylated RPLP1. Additional phosphoproteomics approaches as well as studies to identify the relevant phosphatase are important future objectives.

Our studies focused on the impact of sustained high glucose on ribosome association of RPLP1 in β-cells and islets and its consequences for β-cell function. The extent to which glucotoxic conditions alter RPLP1 ribosome association and translation in other tissues impacted by diabetes complications remains to be determined. Additional work will also be required to understand if other metabolites lead to remodeling of ribosome structure and function. Future studies addressing these questions will help establish the broader physiological relevance of dynamic ribosome heterogeneity.

## RESOURCE AVAILABILITY

### Lead contact

Further information and requests for resources and reagents can be directed to the lead contact, Jean Schaffer

### Materials availability

Plasmids for expression of epitope-tagged and mutated RPLP1 are available upon request to the lead contact.

### Data and code availability

No original code was generated in this study.

Proteomics data generated in this study is deposited in PRIDE (PXD085281).

## ACKNOWLEDGMENTS

We are indebted to members of the Schaffer, Bonner-Weir, and Aguayo-Mazzucato labs for insightful discussions. We thank Jun-ichi Miyazaki for providing MIN6 cells. This work was supported by funding to J.E.S. (NIH R01DK138605, NIH DP1DK119141, Beatson Foundation 2022-005), to ACH (K99DK140526, Mary K Iacocca Senior Postdoctoral Fellow), to JSNA (NSF DGE21240743), to CAM (NIH R01 DK132535), and the Joslin Diabetes Research Center (NIH P30DK036836). The authors also thank the Joslin Islet Isolation Core and the Thermo Fisher Scientific Center for Multiplexed Proteomics at Harvard Medical School.

## AUTHOR CONTRIBUTIONS

ACH and JES conceived and designed the study and analyzed the data. ACH, JSNA, OA, LN, JHL, and KI performed experiments. SBW and CAM analyzed the data and provided intellectual input into the study. ACH and JES wrote the manuscript draft with contributions from JSNA, OA, and LN, and all authors discussed the results, commented on the manuscript, and approved the final version.

## METHODS

### EXPERIMENTAL MODELS AND SUBJECT DETAILS

#### Animals

All experimental procedures and methods were conducted in compliance with institutional guidelines at the Joslin Diabetes Center. All studies involving animals adhered to US Department of Agriculture and National Institutes of Health regulations. Experimental protocols were reviewed, approved, and monitored throughout by the Institutional Animal Care and Use Committee (IACUC) to ensure compliance with institutional animal welfare standards. All animals were housed at 22°C under a 12-h light/dark cycle with access to food and water ad libitum except as noted during experimental procedures. Sprague-Dawley rats were purchased from Taconic Biosciences. NCG mice (NOD-*Prkdc^em26Cd52^IL2rg^em26Cd^*^22^/NjuCrl) were purchased from Charles River (strain 572). Rosa26-floxed STOP-Cas9 knockin mice (B6J.129(B6N)-*Gt(ROSA)26Sor^tm1(CAG-cas9*,-EGFP)Fezh^*/J, strain 026175 on B6J background) and *Ins1^Cre^* (B6(Cg)-*Ins1^tm1.1(cre)Thor^*/J, strain 026801) were purchased from Jackson Laboratories. Rosa26-floxed STOP-Cas9 knock-in mice were crossed with *Ins1^Cre^* mice, with monitoring of genotypes by PCR assay (Transnetyx).

#### Human islets

Human islets from non-diabetic cadaveric donors were obtained from ProdoLabs. Since these are de-identified samples from non-living subjects, these samples were not considered human studies research and did not require additional IRB approval.

### METHOD DETAILS

#### Cell lines

Low-passage MIN6 cells were provided by Jun-ichi Miyazaki (Osaka University, Japan) and were cultured in DMEM containing 4.5 g/L glucose and 4 mM glutamine and supplemented with 15% FBS, 0.1 mM β-mercaptoethanol, 100 U/mL penicillin, and 100 μg/mL streptomycin at 37°C with 5% CO_2_. For analyses of ribosomal proteins, MIN6 cells were treated for 24 h with DMEM containing 5.5 mM or 25 mM glucose prior to analysis of polysomes, proteins, and phospho-proteins. HEK 293T cells from American Type Culture Collection (ATCC CRL-3216) were cultured in DMEM containing 4.5 g/L glucose, supplemented with 10% inactivated FBS and 2 mM L-glutamine, at 37°C with 5% CO_2_.

#### Islets

Islets were isolated from 7 to 8-week-old male Sprague-Dawley rats (Taconic Biosciences) by collagenase digestion followed by density gradient centrifugation by the Joslin Islet Isolation Core.^71^ Handpicked islets were incubated overnight in RPMI 1640 containing 11 mM glucose, 10% FCS, 100 U/mL penicillin, 100 μg/mL streptomycin at 37°C with 5% CO_2_. For analysis of impact of sustained high glucose, rat islets were cultured for 4 days in RPMI 1640 media containing 5.5 mM or 16.7 mM glucose prior to analysis. For lentiviral transduction, freshly isolated islets were transduced immediately and then cultured for 7 days prior to transplantation.

Human islets from cadaveric human donors (Prodo Labs, ages 30-50 years, non-diabetic, see Supplemental Table 2) were incubated overnight in RPMI 1640 containing 5.5 mM glucose, 10% FCS, 100 U/mL penicillin, 100 μg/mL streptomycin at 37°C with 5% CO_2_. For analyses of impact of sustained high glucose, islets were cultured in RPMI 1640 media containing 5.5 mM or 20 mM glucose for 48 h prior to analysis.

#### Sucrose density gradient fractionation

MIN6 cells were treated with 100 μg/mL of cycloheximide for 5 minutes prior to lysis as previously described.^72^ Sucrose gradients (5-50% for large gradients, 15-50% for small gradients) were generated using a BioComp Gradient Master IP and solutions made in sucrose buffer (10 mM Tris [pH 7.2], 60 mM KCl, 10 mM MgCl_2_, 1 mM DTT, and 0.1 mg/mL heparin). Cell lysates were layered onto gradient, centrifuged (Beckman Coulter: SW41T for large gradients [222,200 x *g*, 3 h, 4°C] or TLS55 for small gradients [214,000 x g, 70 min, 4°C]), and polysome and soluble fractions collected using a BR-188 Density Gradient Fractionation System (Brandel). For translation efficiency analysis, RNA was extracted from combined polysome fractions using TRIzol LS (Invitrogen), treated with heparinase (NEB) for 30 minutes at 30° C, and cleaned up with Direct-zol RNA miniprep kit (Zymo Research). For protein analyses, combined polysome fractions were concentrated using Amicon Ultra Centrifugal filters (Millipore Sigma) with 3 kDa cut-off and precipitated by methanol-chloroform method. For immunoblots, dried pellets were resuspended in RIPA buffer (150 mM NaCl, 50 mM Tris pH 8.0, 0.5% sodium deoxycholate, 0.1% SDS, 1% NP40) containing cOmplete EDTA-free Protease Inhibitor Cocktail (Millipore Sigma) and phosphatase inhibitors (Pierce). For proteomics, dried protein pellets were re-suspended in buffer containing 8M Urea, 1% SDS, 200 mM EPPS (pH 8.5), cOmplete EDTA-free Protease Inhibitor Cocktail (Millipore Sigma).

#### Polysome proteomics

Polysome fraction protein (50 µg) was reduced with TCEP, alkylated with iodoacetamide, and further reduced by DTT. Proteins were digested with Lys-C (1:50) overnight at room temperature and trypsin (1:50) for 6 h at 37°C. Peptides were labeled with TMTPro 16plex reagents and desalted using Sep-pak, fractionated into 24 fractions using basic reverse phase HPLC. Samples were analyzed on an Orbitrap Lumos mass spectrometer. MS2 spectra were searched using the COMET algorithm against a mouse Uniprot composite database containing its reversed complement and known contaminants.^73^ Peptide spectral matches were filtered to a 1% FDR using the target-decoy strategy combined with linear discriminant analysis. Proteins were filtered to a <1% FDR and quantified only from peptides with a summed SN threshold >160. Protein intensity was log_2_ transformed, missing values were imputed, and data were normalized such that all samples had the same median abundance.^74^ Linear modeling and moderated t-tests, with adjustment for surrogate variable analysis were performed using limma, as previously described.^75,76^

#### Immunoblotting

MIN6 cells were lysed in RIPA buffer (50 mM Tris [pH 8], 150 mM NaCl, 0.5 % sodium deoxycholate, 1% NP-40, 0.1% SDS) containing cOmplete EDTA-free Protease Inhibitor Cocktail (Millipore Sigma) and protease inhibitors (Pierce), with quantification using Bicinchoninic Acid (BCA) assay (Pierce). Rat and human islets were lysed by sonication in islet lysis buffer (5 mM EDTA, 7 M urea, 2 M thiourea, 100 mM sodium fluoride, 100 mM pyrophosphate, 10 mM orthovanadate, 50 mM PMSF, 1 μg/mL aprotinin (Pierce), and 1% Triton with quantification using Bio-Rad protein assay. Ten to twenty micrograms of protein lysates and immunoprecipitation samples were denatured by boiling in Laemmli sample buffer with β-mercaptoethanol, separated using 10% NuPAGE Bis-Tris gels, transferred to PVDF membranes using iBlot2 (Thermo Fisher), blocked with 5% BSA or 5% non-fat dry milk in TBST, and blotted for the indicated proteins using commercially available antibodies (see table for products and dilutions). HRP-or AlexaFluor-coupled secondary antibodies and chemiluminescent substrates (Bio-Rad or Sigma) were used for detection with a ChemiDoc MP Imaging System (Bio-Rad). Antibodies and dilutions are provided in Supplemental Table 3.

#### Isolation of actively translating ribosomes from rat and human islets

Translating ribosomes were isolated using the AHARIBO RNA system (Immagina Biotech). Briefly, islets were incubated in methionine-free treatment media for 40 minutes, followed by addition of Azidohomoalanine (AHA) for 3 minutes, and sBlock for 5 minutes. Islets were lysed in lysis buffer with sonication, click reaction performed according to the manufacturer’s protocol, and protein eluted using Laemmli buffer and boiling. Input and eluted protein samples were used in immunoblotting to detect total and translating ribosomal proteins, respectively.

#### Lentivirus production and transduction

Short hairpin RNA (shRNA) plasmids (scrambled control and Rplp1-targeting) were obtained from Origene (TL514889). FLAG-tagged lentiviral plasmids expressing Rplp1 wild type (WT) and mutants (S101A and S104A; S101A, S104A) were designed and purchased from VectorBuilder. Lentiviral plasmids and lentiviral packaging mix (Cellecta) were transfected into HEK 293T cells using Lipofectamine (Thermo Fisher), and viral supernatants were collected days 2-4 post-transfection, pooled, concentrated using Lenti-X concentrator (Takara Bio), aliquoted, and stored in −80 °C for transductions. MIN6 cells were transduced with concentrated lentivirus diluted in growth media with 8 µg/ml polybrene (Millipore Sigma). Freshly isolated rat islets (150-200 islets in 100 µL media droplets) were transduced by dropwise addition of 50 μL of 50x concentrated virus, incubation for 5-6 h at 37°C with 5% CO_2_ and subsequent addition of growth media. Analyses were performed 2-3 days (MIN6) or 5 days (islets) following transduction. See Supplemental Table 4 for plasmid sequences.

#### Metabolic labeling of RPLP1 for ribosome association dynamics

Metabolic labeling of pre-existing (old) RPLP1 was performed prior to low vs. high glucose treatment. Newly synthesized (new) RPLP1 was labeled during the final h of low vs. high glucose treatment. MIN6 cells were changed to methionine-free media for 30 minutes, followed by addition of 250 μM AHA (Vector Labs) for 8 h. Cells were then treated with 100 μg/mL of cycloheximide for 5 minutes prior to lysis in ribosome profiling lysis buffer and sucrose density gradient fractionation. Polysome fractions were combined, reduced with 15 mM DTT, alkylated with 20 mM iodoacetamide (Thermo Fisher), quenched with 10 mM DTT, and precipitated with methanol and chloroform. The pellets were resuspended in buffer containing 2% SDS, 50 mM HEPES pH 7.2, 150 mM NaCl, and cOmplete EDTA-free Protease Inhibitor Cocktail (Millipore Sigma) and phosphatase inhibitors (Pierce) with sonication. 1mg protein was incubated with 50 μM TBTA Ligand (Vector Labs), 0.5 mM sodium ascorbate (Sigma Aldrich), 0.5 mM Copper (II) Sulfate (Thermo Fisher), and 50 μM biotin-alkyne reagent (Vector Labs) for 2 h at room temperature. Proteins were precipitated with methanol and chloroform, resuspended with 2% SDS, 5 mM DTT containing protease and phosphatase inhibitors, and diluted in RIPA containing protease and phosphatase inhibitors. FLAG-RPLP1 was immunoprecipitated, separated by SDS-PAGE, transferred to PVDF membrane, analyzed by immunoblotting for FLAG and binding of streptavidin-HRP, and quantified with normalization to immunoblotted FLAG-RPLP1 in input samples.

#### Immunoprecipitation

MIN6 cells were lysed in TNET buffer (50 mM Tris [pH 8.0], 150 mM NaCl, 2 mM EDTA, 1% Triton X100) containing cOmplete EDTA-free Protease Inhibitor Cocktail (Millipore Sigma) and Phosphatase Inhibitors EDTA-free Mini Tablets (Pierce A32957), triturated using 27-gauge needles, and cleared by centrifugation (20,000xg, 10 min, 4°C). Rat islets were lysed in RIPA buffer containing cOmplete EDTA-free Protease Inhibitor Cocktail (Millipore Sigma), Phosphatase Inhibitor Cocktail 3 (Millipore Sigma), Phosphatase Inhibitor EDTA-free Mini Tablets (Pierce), and 2 mM EDTA; triturated using 27-gauge needles; sonicated; and cleared by centrifugation (20,000 x g, 10 min 4°C). Precipitated polysomes were resuspended in RIPA buffer for immunoprecipitation. Samples were pre-cleared with α-HA magnetic beads (Millipore Sigma SAE0197, nutated 15 min, room temp). Lysates were incubated with α-FLAG magnetic beads (Millipore Sigma M8823) (nutated 1 h, room temp), and beads were washed 3 times with modified TNET high salt buffer (50 mM Tris [pH 8.0], 500 mM NaCl, 2 mM EDTA, 1% Triton X100) containing protease and phosphatase inhibitors and then PBS. Immunoprecipitates were recovered by boiling in 2X Laemmli Sample Buffer (Bio-Rad).

#### Detection of phosphorylated proteins using Pro-Q Diamond

Following protein separation on NuPAGE 10% Bis-Tris gels, gels were fixed with 10% acetic acid/50% methanol, washed 3 times in distilled water (10 min each), incubated with Pro-Q Diamond phospho-protein stain for 1.5 h, destained 3 times in Pro-Q Diamond destain solution (30 min each), and washed in distilled water. Gels were imaged in ChemiDoc MP Imaging System (Bio-Rad) using Pro-Q Diamond setting (excitation 555 nm, emission 580 nm). For dephosphorylated samples, lysates were treated with 10 U FastAP Thermosensitive Alkaline Phosphatase (rotating, 1 h, room temp) before pre-clearing with anti-HA magnetic beads and immunoprecipitations were performed without phosphatase inhibitors. Parallel immunoblot analysis was performed on immunoprecipitates, with densitometric quantification of Pro-Q Diamond stain signal relative to immunodetected signal for each immunoprecipitate.

#### Detection of nascent proteins using OPP

Cells were incubated for 1 h in media containing 20 μM *O*-propargyl-puromycin (OPP; Click Chemistry Tools) and click reaction for in-gel detection of nascent peptides was performed as previously described,^20^ and proteins were separated on NuPAGE 10% Bis-Tris gels. Following fixation with 10% acetic acid, 50% methanol, nascent peptides were detected by in-gel imaging at 647 nm, and total protein was detected by Epi-far red imaging after staining with 0.1% Coomassie brilliant blue (Bio-Rad ChemiDoc MP Imaging System).

#### Isolation of ribosome-associated mRNA from islets

Approximately 500 islets were treated with 100 μg/mL of cycloheximide for 5 minutes, washed with ice-cold PBS, and lysed with ribosome profiling lysis buffer. Two-thirds of each sample was centrifuged through a 1M sucrose cushion (Beckman Coulter: TLA100.2, 436,000 x *g*, 1 h, 4°C) to collect ribosome-associated mRNA in pellet, and the remaining one-third was collected for total RNA isolation. RNA isolations were done by phase separation method, using TRIzol (Invitrogen) and 1-bromo-3-chloropropane (Millipore Sigma).

#### mRNA quantitative reverse transcription (RT) PCR

RNA was recovered from sucrose density gradient fractions using DirectZol RNA extraction kit (Zymo Research). RNA was treated for 20 minutes at room temperature with 600 U/mL heparinase (NEB P0735S). 500ng to 1 μg RNA was reverse transcribed using iScript cDNA synthesis kit (Bio-Rad) or SuperScript (Thermo Fisher). Quantitative RT PCR was performed using SsoAdvanced Universal SYBR Green Supermix (Bio-Rad) and gene-specific primers (Supplemental Table 4). Relative mRNA abundance was calculated based on the ΔΔCt method relative to 18S rRNA.

#### Glucose stimulated insulin secretion (GSIS)

Static incubation GSIS was performed as previously described.^20^ Briefly, MIN6 cells (10^5^ cells/35-mm well, assayed in duplicate) or 10-12 islets (∼150 μm diameter, assayed in triplicate) were washed and incubated with Krebs-Ringer bicarbonate HEPES buffer (KRBH) containing 2.8 mM glucose for 1 h. This was followed by successive incubation with fresh KRBH-2.8 mM glucose and KRBH-high glucose (16.8 mM for MIN6, 16.7 mM for islets), each for 1 h. Supernatant and cells were collected for insulin measurement using Ultra Sensitive Mouse Insulin ELISA (Crystal Chem). Insulin was normalized to cell number for MIN6 cells or DNA content (Cyquant Proliferation kit, Thermo Fisher) for islets.

#### Intraperitoneal Glucose Tolerance Test (IPGTT)

Mice were fasted for 6 h with ad libitum access to water prior to initial blood collection from tail vein and quantification by glucometer. Animals with fasting blood glucose < 350 mg/ dl received intraperitoneal injection of 2g/kg body weight of 10% glucose solution. Blood was collected at before (0) and 15, 30, 60, and 120 minutes after glucose injection for glucometer analysis.

#### Rat Islet transplantation

NCG 8-week-old male mice were rendered diabetic (fed blood glucose between 380 and 595 mg/dl at day 7 post-injection) by one-time IP injection of 140 mg/kg streptozotocin (Millipore Sigma). Rat islets transduced with RPLP1-targeting and control shRNAs were cultured in the RPMI 1640 (11mM glucose) media for 7 days prior to transplantation. 400 IEQ were transplanted under the left kidney capsule by using a Hamilton syringe (Fisher Scientific #14-824-40) attached to polyethylene tubing P-50 (Fisher Scientific 14-170-12B).^77^ Animals were monitored for 2 weeks prior to IPGTT by morning fed glucose and only animals that achieved blood glucose < 350 mg/dl were used in IPGTT study at day 14 post-transplant.

#### AAV8-gRNA editing in pancreatic β-cells

AAV vector plasmids were synthesized by VectorBuilder Inc. and contained two guide RPLP1-targeting or scramble gRNAs within the 5′ and 3′ ITRs (U6 promoter upstream of the first guide RNA and macU6 promoter upstream of the second guide RNA) and a WPRE element within the ITRs (sequences in Supplemental Table 4). Large-scale AAV8 preparations (>10^13^ GC/ml) were generated by VectorBuilder Inc. At 4-5 weeks of age, Ins1-Cre^+/-^, cCAS9^FL/FL^ female mice were injected IP with AAV8 (4 × 10^12^ GC in 100 µl saline). Morning fed blood glucose was monitored two and three weeks post-injection. IPGTT was performed at 4 weeks post injection.

#### Immunohistochemistry

Animals were euthanized and pancreases removed following IPGTT. Tissues were fixed in 4% paraformaldehyde (2 h) and processed for paraffin embedding. Sections were immunostained for insulin and RPLP1 (Supplemental Table 3). After deparaffinization, slides underwent antigen retrieval by microwaving in 10 mM Tris pH 9, 1 mM EDTA, 0.05% Tween-20. Slides were successively blocked with avidin and biotin (Vector Laboratories) according to manufacturer’s recommendations, followed by normal donkey serum (1:50) in PBS. RPLP1 staining (Proteintech, 1:250) was at 4°C overnight, goat α-rabbit biotin staining (Jackson ImmunoResearch, 1:200) was for 1 h at room temperature (RT), streptavidin-AlexaFluor 594 (Jackson ImmunoResearch, 1:200) was for 1 h at RT, guinea pig α-insulin (Thermo Fisher,1:50) for 1.5 h at RT, and goat α-guinea pig AlexaFluor 647 (Jackson ImmunoResearch,1:200) was for 1 h RT, with all incubations and washes in PBS with 2% sheep serum (Thermo Fisher). Coverslips were mounted with VectaShield Antifade mounting media with DAPI (Vector Labs). Images were acquired using a Zeiss Observer 7 equipped with a monochromatic camera (Axiocam 705 mono) and a true-color camera (Axiocam 305 color), using Colibri LED IIluminators that include the following excitation channels: 385 nm for DAPI, 475 nm for GFP, 555 nm DS red (AlexaFluor 594), 630 nm for Cy5, (AlexaFluor 647). Images were captured with the Plan-Apochromat 20x/0.8 Ph2 or Plan-Apochromat 40x/0.95 Corr and Zeiss Zen v. 3.10.103.07000 software. Fiji/ImageJ version 2.16.0/1.54p was used for quantification from 20X images. RPLP1 staining intensity was quantified in regions of interest (ROI) defined by insulin staining for n = 3 animals per experimental group, with a minimum of 10 islets/animal.

